# *Escherichia coli* siphovirus DH23 features unusual C12-C13-C6 symmetry of the neck complex and the lateral tail fibers branching from the central tail fiber

**DOI:** 10.64898/2026.11.25.768851

**Authors:** M.A. Bubenchikov, P.A. Ivanov, O.S. Sokolova, A.V. Moiseenko, A.V. Letarov

## Abstract

Bacteriophages capable of propagating in host cells that have entered or are approaching the early stationary phase of growth (eSPI phages) are promising candidates for phage therapy as they may efficiently destroy slowly growing pathogen populations *in vivo*. Among the *Escherichia coli* phages, the eSPI phenotype is rare and the diversity of such phages is limited. Siphoviruses of the *Dhillonvirus* genus appear to offer an attractive platform for implementing host range management systems. We investigated the structure of the eSPI Dhillonvirus DH23 using cryo-electron microscopy and single-particle analysis and revealed that its adaptor protein connecting the dodecameric head portal to the hexamer ring of the tail termination protein forms a tridecameric (C13) ring rather than a dodecameric ring as in most known bacteriophages. Despite the presence of the putative lateral tail fiber (LTF) protein gene in the DH23 genome, the tail tip 3D density map showed no evidence of LTF attachment. Examination of 2D projection images of the flexible distal part of the central tail fiber (CTF) revealed a massive density from which the LTFs emanate, forming a branched CTF-LTF structure that was not seen in coliphages previously.

## Introduction

Bacterial viruses, known as bacteriophages (phages), are currently considered promising antibacterial agents capable of controlling even pathogenic strains resistant to most or all antibiotics available for clinical use (1, 2). Pathogenic *Escherichia coli* has been identified by the WHO as one of the leading bacterial species contributing to global mortality associated with drug-resistant bacterial infections (3). Although most tailed bacteriophages propagate efficiently in actively growing logarithmic-phase laboratory cultures, their multiplication is severely inhibited as the culture approaches or enters early stationary phase (4). Interestingly, some *E. coli* bacteriophages are able to effectively infect and lyse bacterial cultures in the early stationary phase (eSPI phages) or even at later stages of bacterial growth (5–7).

These phages are considered potentially more effective in phage therapy (PT) than “conventional” phages. However, the limited diversity of known eSPI bacteriophages hinders the development of PT approaches based exclusively on natural isolates of these viruses. Therefore, the eSPI phage platforms suitable for implementation of artificial host range modification systems retargeting them to specific pathogenic strains are of particular interest. Phage infection of *E. coli* is significantly limited by the non-specific protection of the bacterial cell surface by the O-antigens (8). Bacteriophages exploit multiple strategies to penetrate this barrier (8). In siphoviruses, lateral tail fibers (LTF) are often essential for the interaction with the O polysaccharide (OPS) (8–10) enabling subsequent interaction of the central tail fiber (CTF) with more conserved secondary receptors such as the outer membrane proteins (11–13). Among the eSPI coliphages characterized to date, the majority are siphoviruses, belonging either to the family Drexelviridae (related to the phage T1) or to the genus *Dhillonvirus* (currently not included in any family) (7). Interestingly, T1-related phages do not contain genes encoding putative LTFs in their genomes and most of Drexelviruses that infect the rough *E. coli* K-12 MG1655 strain are strongly restricted by the O16 O-antigen in the restored wbbl+ derivative (14). The representatives of *Dhillonvirus* genus, in contrast, contain the potential LTF gene and some of these viruses have been shown to infect the O16 – restored *E. coli* MG1655 derivative (15). A reconstruction of the Dhillonvirus Bas18 (16) — which is closely related to DH23, with which it shares more than 95% amino acid sequence identity across most structural proteins — was recently published, based on cryo-electron microscopy (cryo-EM). Interestingly, despite the relatively high resolution of the tail tip complex of this phage no evidence of LTF attachment could be observed in this reconstruction. In addition, the Bas18 bacteriophage contains a poorly resolved collar-like density surrounding the neck region which is an unusual feature for siphoviruses.

Previously we described the Dhillonvirus DH23 featuring the eSPI phenotype similar to that of the classical coliphage T7 with eSPI (17). The genome of this phage contains a gene encoding the putative LTF protein gp30. Here we performed cryo-EM of the DH23 virions and focused on resolving the collar-like density and determining the location of LTF. We show that the apparent collar is formed by side domains of the adaptor protein located between the bacteriophage head portal protein and the tail completion protein. Unexpectedly, the DH23 adaptor protein forms a C13-symmetric ring, whereas the adaptor rings of all previously characterized tailed bacteriophages are, to our knowledge, C12-symmetric. The DH23 tail tip was depleted of the LTFs as in the Bas18 phage. However, the examination of the distal CTF region, which was absent from the high-resolution density map due to the CTF flexibility, allowed us to visualize the LTFs branching from the CTF.

## Materials and methods

### Bacteriophage and bacterial strains and their propagation

Phage DH23 was isolated and characterized as described by us previously (17). Its genome was sequenced and deposited in GenBank under accession number LC891837. The *E. coli* MG1655 strain was obtained from our laboratory collection. The LB medium (Trypton 10g, Yeast extract 5g, NaCl 10g, deionized water up to 1L) Bacto-agar per 1L for the plates or 6g for the soft agar used for the double-layer bacteriophage titration.

For growing the DH23 lysate the overnight host strain culture was grown up at 37°C and 250 rpm agitation. This culture was then diluted 1:100 in LB medium in the Erlenmeyer flask and propagated for 6 hours under the same conditions. The culture was then inoculated with approximately 10^7^ PFU of the DH23 phage per 1 mL. The incubation was continued overnight. The phage lysate was clarified by centrifugation at 5000 g for 10 min at ambient temperature and filtered through the syringe membrane filter of 0.22 μm pore size. The lysate titer was determined by conventional double-layer plaque assay. The typical titer ranged from 2–7 × 10^11^ PFU mL^-1^

### Phage purification for cryo-electron microscopy

DH23 sample preparation was performed using a slightly modified Ackermann’s protocol (18). Briefly the phage particles from 15 mL of phage lysate were pelleted by centrifugation at 50,000 g for 1 h in a fixed-angle rotor. The supernatant was carefully removed, and the pellet was gently resuspended in the original volume of 0.1 M NH_4_Cl solution at pH 7.5. The centrifugation and resuspension steps were repeated twice, and the phage was finally resuspended in 1 mL of the same buffer. Phage suspension was left overnight at 4°C and the aggregates were removed by centrifugation at 10,000g for 10 min in a benchtop microcentrifuge.

The phage purity was assessed by negative-stain transmission electron microscopy performed as it described previously (17).

### Cryo-EM data collection

Purified DH23 samples were applied to Quantifoil R1.2/1.3 copper grids and plunge-frozen in liquid ethane using Vitrobot Mark IV. The sample was imaged using TOTEM300s cryo-electron microscope (Shuimu Biosciences, PRC), equipped with Gatan K3 detector. Movies were collected in counting mode with an electron dose of 50 e/Å2, at magnification corresponding to a pixel size of 0.85 Å. A total of 5,424 movies were collected.

### Single Particle Analysis of DH23

All image processing was fully conducted in CryoSPARC v4.7.1 (19). Movies were subjected to patch-based motion correction and patch-based Contrast Transfer Function estimation. The dataset was curated by removing low-quality micrographs, based on measured relative ice thickness and the maximum Contrast Transfer Function fit spatial frequency. A total of 4,986 micrographs were used for particle picking.

#### Tail tip reconstruction

To investigate the tail tip of phage DH23, we used Filament Tracer with a filament diameter of 120 Å and a large separation distance 96 Å to increase the proportion of tail tip particles relative to tail tube particles in Filament Tracer output. After 2D classification, 1,749 tail tip particles were used to train Topaz (20) picker. The resulting particles were also subjected to 2D classification and used as input data for a second Topaz distinct run. This step was repeated once more, and 11,635 particles were passed through Ab-Initio Refinement, Homogeneous Refinement and Local Refinement with C3 symmetry. To further increase resolution, Local and Global Contrast Transfer Function Refinement followed by Reference-Based Motion Correction were conducted. After final Homogeneous and Local Refinements with C3 symmetry, the tail tip reconstruction with a resolution of 3.38 Å was obtained.

After obtaining the consensus tail tip reconstruction, we adjusted the coordinates of aligned particles using Volume Alignment tools to center the box on the distal tip region. Then 2D classification was applied to discover heterogeneity in the distal tail tip region.

#### Neck reconstruction

Manual particle picking with a diameter of 850 Å was used to generate templates for automatic picking of capsid projections. After 2D classification, a subset of 52,575 particles was selected for Ab-Initio Reconstruction with icosahedral symmetry imposed. The resulting volume was used as a reference for Homogeneous Refinement with icosahedral symmetry imposed. The final consensus volume was resolved at 3.23 Å according to the gold standard Fourier Shell Correlation 0.143 criterion (GSFSC).

To identify the unique portal vertex, capsid particles were symmetry-expanded using icosahedral symmetry and subjected to 3D classification focused on the vertex region. The class containing tail density was subjected to Local Refinement with C12 and C6 symmetry imposed, focusing on the portal and distal neck regions, respectively. The resulting reconstructions had resolutions of 3.01 Å for the portal region and 3.05 Å for the distal neck region. However, the collar region remained poorly resolved in both volumes, necessitating an asymmetric C1 reconstruction.

To obtain the C1 reconstruction of the portal-capsid region the portal region was 3D classified to identify distinct rotational states of the portal relative to the capsid. These states were rotated in increments of 72° to align them in a common orientation. Then this dataset was subjected to Local Refinement without imposing symmetry. Although the capsid and portal were both aligned, the collar region remained poorly resolved. 3D classification focused on the collar region was used to investigate its density. The classification yielded two classes corresponding to two distinct rotational states. The asymmetric reconstruction clearly revealed that the adaptor ring is C13-symmetric. The collar region was then subjected to Local Refinement with C13 symmetry imposed. Local and Global Contrast Transfer Function Refinement jobs were applied to further improve the resolution. The final reconstruction of the collar region exhibited an unusual C13 symmetry and had a resolution of 3.15 Å.

### Model building

The sequence of the tail adaptor protein was previously published (GenBank accession number LC891837). The initial model was built based on this sequence using the local implementation of Alphafold 2 in ColabFold v1.5.5 (21, 22). Further refinement was performed using UCSF ChimeraX v.1.11.1 with the ISOLDE bundle v1.11.1 (23, 24). The final model was refined against the density map using *phenix*.*real_space_refine* from PHENIX suite v1.21.2-5419 (25). All figures showing the density maps and atomic models were prepared using UCSF ChimeraX v.1.11.1.

## Results

LTFs are usually invisible in cryo-EM reconstructions because of their flexibility. The DH23 genome includes a gene that encodes gp30, a putative LTF protein (17). To detect the potentially flexible region in this protein, we analyzed its amino acid sequence. The HHpred search revealed strong similarities (probability >97%) between the N-terminal region (residues 100– 110) and the N-terminal regions of the pyocin F2 side tail fibers (PDB ID: 25VE, chain C) and the pyocin F7 N-terminal domains (PDB ID: 9YG8, chain J), as well as to the other phage fibers and tail spikes.

The C-terminal 117 amino acids showed strong similarity (probability >97%) to the C-terminal autocleavable chaperone domain of the LtfA protein from the phage DT57C. This region also contains a conserved motif associated with a potential cleavage site, TS^XXXX(R/K) (9) , suggesting that the mature gp30 protein may terminate at residue S358. The AlphaFold 3 structure prediction suggests that the DH23 LTF forms a straight rod with a globular domain at its C-terminus, potentially involved in receptor binding. A potentially flexible region is predicted between this domain and the coiled-coil region (Fig. 1).

**Fig. 1:**
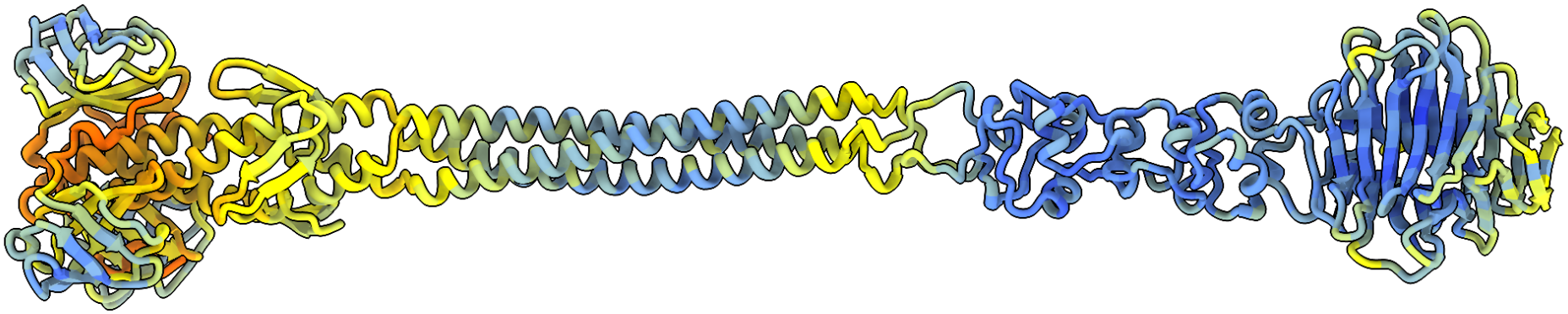
Predicted LTF structure (gp30 trimer) colored by Alphafold’s pLDDT. The C-terminal chaperon domain is removed.

Although this protein is considerably shorter than LTF proteins from other siphoviruses (for example, 1,264 residues in the processed LTF protein of phage T5 and 774 residues in the Stf protein of phage Ur-λ), its N-terminal region appears typical of bacteriophage fibers and should be easily detectable if attached to the tail-tip complex.

We next obtained cryo-EM data for whole phage DH23 (Fig. 2 A) and generated the density maps of the tail tip and neck regions. The tail tip complex appeared nearly identical to the recently reported structural model of bacteriophage Bas18 (17) , consistent with the high sequence similarity between the structural proteins of these two phages. As in phage Bas18, approximately 820 N-terminal residues of CTF protein gp26 form a distal part of the tail tip cone. No LTFs attachment was observed in 3D reconstruction.

**Fig. 2:**
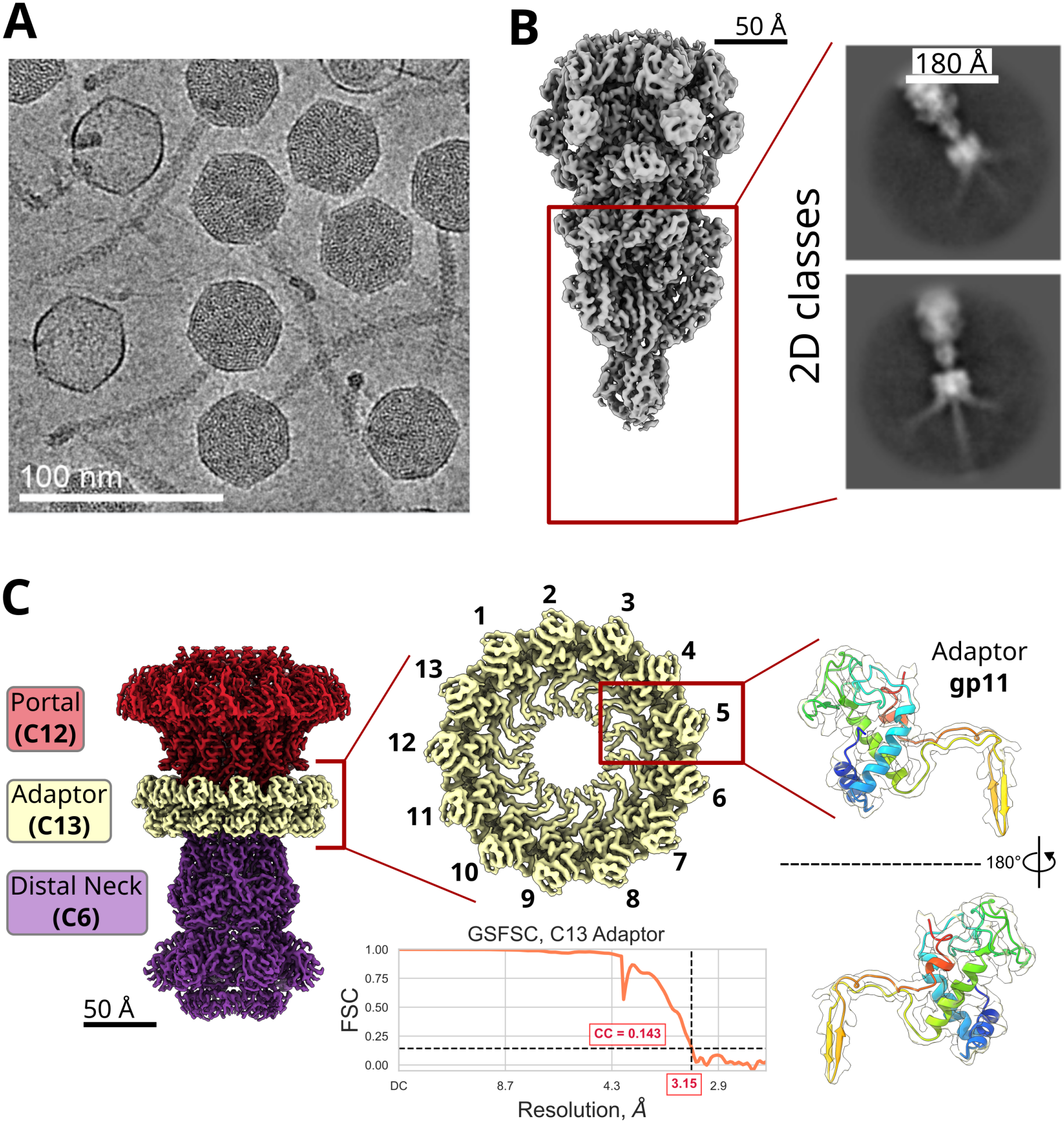
Cryo-EM analysis of DH23 bacteriophage parts. A – raw image of virions. B – tip volume and 2D classes of the distal tail tip region with a clearly observable large complex with branching fibers. C – analysis of DH23 neck. Left - There are three types of symmetries – C12 for the portal, C6 for the distal neck, and unusual C13 symmetry for adaptor region. Middle - 13 copies of adaptor protein can be easily distinguished. The map resolution is sufficient for atomic model building. Right - The adaptor gp11 atomic model is colored in rainbow spectrum (from blue N- to red C-terminus).

To resolve a flexible distal tip region, we employed 2D classification after recentering the tail tip particles. The resulting 2D classes clearly revealed a massive complex assembled on the CTF with fiber-like densities extending on either side of the CTF. The longer co-axial density of the CTF is also visible (Fig. 2 B).

We interpret these densities as gp30 complexes attached to the CTF, forming the branched structure. Recently a similar tripod-like structure has been reported for the F2-type pyocin from *P. aeruginosa* ATCC 15442 (26). In the F2 pyocin the CTF and three LTFs form a more rigid, pyramid-shaped complex which allowed to build a high resolution structural model. In the phage DH23 the components of the branching point appear to be more flexible, currently preventing us from more detailed reconstruction of this structure. We cannot reliably determine the symmetry or the number of the LTFs present although 2D classes obtained can accommodate three or four fibers. Nevertheless, we clearly demonstrate the attachment of the side fibers to the needle-like part of the CTF. Although the resolution of the LTFs is by far not sufficient to directly identify the protein(s) from which they are formed, the relatively small (44,682 bp) genome of the phage DH23 does not encode any potential candidates besides gp30.

The 3D reconstruction of the bacteriophage DH23 neck region confirmed the presence of the collar-like density, similar to that observed in the phage Bas18 (16). To obtain consistent alignment in the collar region, we generated a C1 reconstruction of the portal-capsid complex and used it to determine the symmetry of the collar. The asymmetric reconstruction clearly revealed that the adaptor ring is C13-symmetric. Then the particle set was subjected to C13 Local Refinement in the collar region. As a result, we obtained a well-resolved density map of C13 adaptor ring, suitable for atomic model building (Fig. 2 C, middle). The adaptor protein is encoded by gene gp11, allowing us to build an atomic model for residues 11-162 of the 176-residue sequence (Fig. 2 C, right). However, investigating the mechanisms that compensate for the symmetry mismatch will require full asymmetric reconstruction of the neck region and will be addressed later.

## Discussion

The overall molecular architecture of the siphovirus tails was long believed to be highly conserved (reviewed in (27)). Recent advances in cryo-EM technology helped to discover the structural features in some viruses significantly deviating from this paradigm. For example, a recent reconstruction of the LTFs of the bacteriophage Ur-λ revealed that in contrast to most siphoviruses having three or six lateral adsorption devices (which is consistent with the C6 symmetry of the tail tip to which these structures are typically attached) the Ur-λ phage carries four LTFs (13). Here we report the first evidence of LTFs attachment to the siphovirus CTF to form a fork-like structure which, up to our knowledge, has not previously been observed in viable bacteriophages capable of delivering the genomic DNA into the host cell.

The observed unusual arrangement of the tail fibers of the DH23 phage raises a question about its compatibility with the established mechanism of action of long, noncontractile phage tails. As it was recently demonstrated for the phages T5 (11) and λ (12, 13), binding the CTF to the receptor induces its bending while bringing the tail tip closer to the outer membrane surface. The presence of the crown of the LTFs mounted around the CTF needle may sterically hinder such binding. Notably, the adsorption curves of the DH23 phage (17) suggest a significant time gap between the initial reversible adsorption of the phage and initiation of the DNA release. This prolonged reversible adsorption phase may reflect the requirement for LTFs dissociation to allow the CTF structural transition. Further experiments are needed to test this hypothesis.

Another surprising feature of phage DH23 and, potentially, other *Dhillonvirus* phages (14, 16) is the unusual C13 symmetry of the gp11 adaptor protein ring. The adaptor connects C12-symmetric portal ring to C6-symmetric tail completion protein. Up to our knowledge, adaptors in all previously characterized phages form either dodecameric rings or multiple rings when more than one adaptor protein is present. The C13 symmetry is not compatible with the neighboring C12 and C6 structures, suggesting that this mismatch must be accommodated by the neck complex specifically. The precise mechanisms underlying the assembly and stabilization of this asymmetric complex remain to be investigated. We currently have no consistent hypotheses regarding the potential ecological advantages that this unusual neck architecture may confer on these phages.

Experimental and bioinformatic evidence suggest that in the *Dhillonvirus* phages the recognition of the terminal protein receptor required for the genome release is mediated by the small dimeric receptor-binding proteins attached to the end of the CTF trimer (14, 28); in DH23 these would be the proteins gp27 and gp28. The LTF function seems to be dispensable for the infection of the rough host strains as observed in many other *E. coli* siphoviruses (reviewed in (8)). The function of the LTF may be connected to the O-antigen penetration which is only essential on smooth (O-antigen-producing) hosts. This trait potentially allows to separately engineer phage DH23 and other dhillonviruses to alter their specificity to different types of the O-polysaccharides and to different outer membrane proteins. Such host-range modification strategy may facilitate the harnessing of these eSPI phages as platforms for the routing generation of the modified bacteriophages for PT of *E. coli* infections.

## Acknowledgments

We are grateful to Dr. E.E. Kulikov and Dr. A. K. Golomidova from RC Biotechnology RAS, Moscow Russia, for their technical assistance in the bacteriophage purification.

We thank Shuimu BioScineces for Cryo-EM facility access and technical support during data collection.

The research was supported by Russian Science Foundation (24-44-02003).

